# *S*-nitrosylation of the CLIP-protease, Persephone, Regulates *Drosophila* innate immunity

**DOI:** 10.64898/2026.01.23.701332

**Authors:** Rafael A. Homem, Krieng Kanchanawatee, Yuan Li, Tao-Ho Chang, Jiayue Zheng, David J. Finnegan, Gary J. Loake

## Abstract

Nitric oxide (NO) modulates innate immunity, but its molecular targets in *Drosophila melanogaster* are largely undefined. A key mechanism of NO signalling is *S*-nitrosylation, the modification of cysteine thiols. The homeostasis of *S*-nitrosylation is maintained by *S*-nitrosoglutathione reductase (Gsnor), encoded by the *fdh* gene in *D. melanogaster*. As reduced Gsnor activity enhances pathogen sensitivity in plants, we investigated its role in *D. melanogaster*. Here, we show that flies lacking *fdh* exhibit increased susceptibility to the fungus *Beauveria bassiana* and the bacterium *Staphylococcus aureus*, pathogens combatted by the Toll pathway. This immune deficiency correlates with impaired Toll-dependent antimicrobial peptide expression. We demonstrate that the Toll pathway protease Persephone (Psh) is *S*-nitrosylated *in vivo* and that loss of Gsnor prevents its proteolytic activation following infection. We propose a model where Gsnor-mediated regulation of NO is essential for Toll activation, preventing excessive *S*-nitrosylation of Psh and ensuring a robust immune response.

## Introduction

The innate immune system is the evolutionarily ancient, first line of defence against invading pathogens, and its core mechanisms are conserved across all metazoan life^1^. The fruit fly, *Drosophila melanogaster*, has long been a genetic model for dissecting the molecular architecture of innate immunity^2^. Its relative simplicity, characterized by the absence of a confounding adaptive immune system, and the profound evolutionary conservation of its core signalling pathways have yielded foundational insights into mammalian immunity, most notably the discovery of Toll-like receptors (TLRs) and their role in pathogen recognition^3^.

In *D. melanogaster*, host defence against systemic infection is orchestrated primarily by two major humoral signalling cascades: the Toll pathway and the Immune Deficiency (IMD) pathway^4^. These pathways exhibit a remarkable degree of specificity, with the Toll pathway mounting a defence predominantly against fungi and Gram-positive bacteria, while the IMD pathway is activated mainly in response to Gram-negative bacteria^5^. Activation of these cascades culminates in the nuclear translocation of distinct NF-κB transcription factors, Dif and Dorsal for the Toll pathway, and Relish (rel) for the IMD pathway, which in turn drive the expression of a battery of effector genes, including those encoding potent antimicrobial peptides (AMPs) such as Drosomycin (Drs) and Metchnikowin (Mtk)^6^.

The activation of the Toll pathway is a sophisticated process initiated by two mechanistically distinct upstream branches. The canonical branch involves the direct recognition of pathogen-associated molecular patterns (PAMPs), such as Lys-type peptidoglycan from Gram-positive bacteria or β-glucan from fungi, by circulating pattern recognition receptors (PRRs) like PGRP-SA and GNBP3. This recognition event triggers a self-amplifying extracellular serine protease cascade that ultimately leads to the cleavage and activation of the cytokine-like ligand, Spätzle (Spz), which then binds to the Toll receptor^7^. In parallel, a second, non-canonical branch functions as a “danger-sensing” system. This pathway is not triggered by static molecular patterns but rather by the functional activity of virulence factors, specifically the proteases secreted by invading microbes. At the heart of this danger-sensing module is Persephone (Psh), a CLIP-domain serine protease that circulates as an inactive zymogen. The pro-Persephone contains a unique “bait” region that is susceptible to cleavage by a broad range of microbial proteases^8^. This initial cleavage licenses pro-Psh for subsequent maturation by endogenous host proteases, unleashing its active form and initiating the downstream cascade that converges on Spz processing^8, 9, 10^. This dual-input architecture allows the Toll pathway to respond not only to the presence of microbes but also to their pathogenic actions.

Superimposed upon these well-defined signalling networks are layers of regulation mediated by pleiotropic signalling molecules. One such molecule is nitric oxide (NO), a highly reactive and diffusible gas radical that functions as a key signalling intermediate in a vast array of physiological processes, from neurotransmission to vasodilation and immunity^11, 12^. A primary mechanism through which NO exerts its biological effects is *S*-nitrosylation, the covalent addition of a NO moiety to the thiol group of a reactive cysteine residue, forming an *S*-nitrosothiol (SNO). This reversible, redox-based post-translational modification acts as a molecular switch, altering protein activity, localization, and stability^13^. Cellular homeostasis of NO and, by extension, global levels of protein *S*-nitrosylation are tightly controlled by the enzyme *S*-nitrosoglutathione reductase (Gsnor). Gsnor catalyzes the metabolism of *S*-nitrosoglutathione (GSNO), the most abundant low-molecular-weight SNO and a major biological reservoir of NO bioactivity. Consequently, mutants with no GSNOR activity provide a powerful tool to investigate the physiological consequences of elevated global *S*-nitrosylation^14, 15^.

While NO has been implicated in *D. melanogaster* immunity^16^, its specific molecular targets and precise regulatory functions, particularly within the complex proteolytic cascades of the Toll pathway, have remained largely undefined. Work in other organisms, such as plants, has revealed that loss of *Gsnor* can paradoxically either enhance or compromise disease resistance, suggesting its role is highly context-dependent^14, 17^. This ambiguity highlights a significant gap in our understanding of how redox signalling is integrated with innate immune activation. Here, we hypothesize that Gsnor-mediated control of NO homeostasis serves as a critical regulatory checkpoint for the *D. melanogaster* Toll pathway. We propose that under conditions of *Gsnor* deficiency, excessive *S*-nitrosylation directly targets and inhibits a key component of the Toll-activating protease cascade. This study sought to identify this molecular target, elucidate the mechanism of inhibition, and explore the broader physiological and evolutionary implications of this novel regulatory axis.

## Results

### Loss of GSNOR Function Compromises Immunity to Toll-Dependent Pathogens

To investigate whether *S*-nitrosylation has a role in the immune response of *D. melanogaster* adults, we reduced Gsnor activity using mutations in the formaldehyde dehydrogenase gene *fdh* (CG6958) that codes for the enzyme with Gsnor activity^18, 19^. At the start of this investigation there were no null alleles of *fdh* available so we generated flies heterozygous for deficiencies (deletions) Df(3R)Exel7305 (Df7305) and Df(3R)Exel7306 (Df7306)^20^ that have breakpoints that overlap within *fdh* (Fig. 1A). These flies have no intact copy of *fdh* (Fig. 1B) and have reduced Gsnor activity that is restored if Gsnor is expressed from an *fdh* transgene using the GAL4-UAS system^21^ (Fig. 1C). Females trans-heterozygous for these overlapping *fdh* deletions have increased levels of nitrite, a proxy NO (Fig. 1D) and also show reduced survival as compared with wild-type (Oregon R) after being fed a 5% sucrose solution containing 5mM sodium nitroprusside (SNP) a phenotype that is also rescued by expression of Gsnor from an *fdh* transgene (Fig. 1E). We have since used CRISPR/Cas9^22^ to replace the coding sequence of *fdh* with the 3xP3-DsRed reporter^23^ resulting in the knockout of the gene. Three independent lines were generated, *fdh^cp1^, fdh^cp2^,* and *fdh^cp3^* (Fig. S1A). Flies homozygous for these alleles have no intact copy of *fdh* (Fig. S1B), have no detectable Gsnor activity (Fig. S1C) and are sensitive to SNP (Fig. S1D).

**Figure 1.**
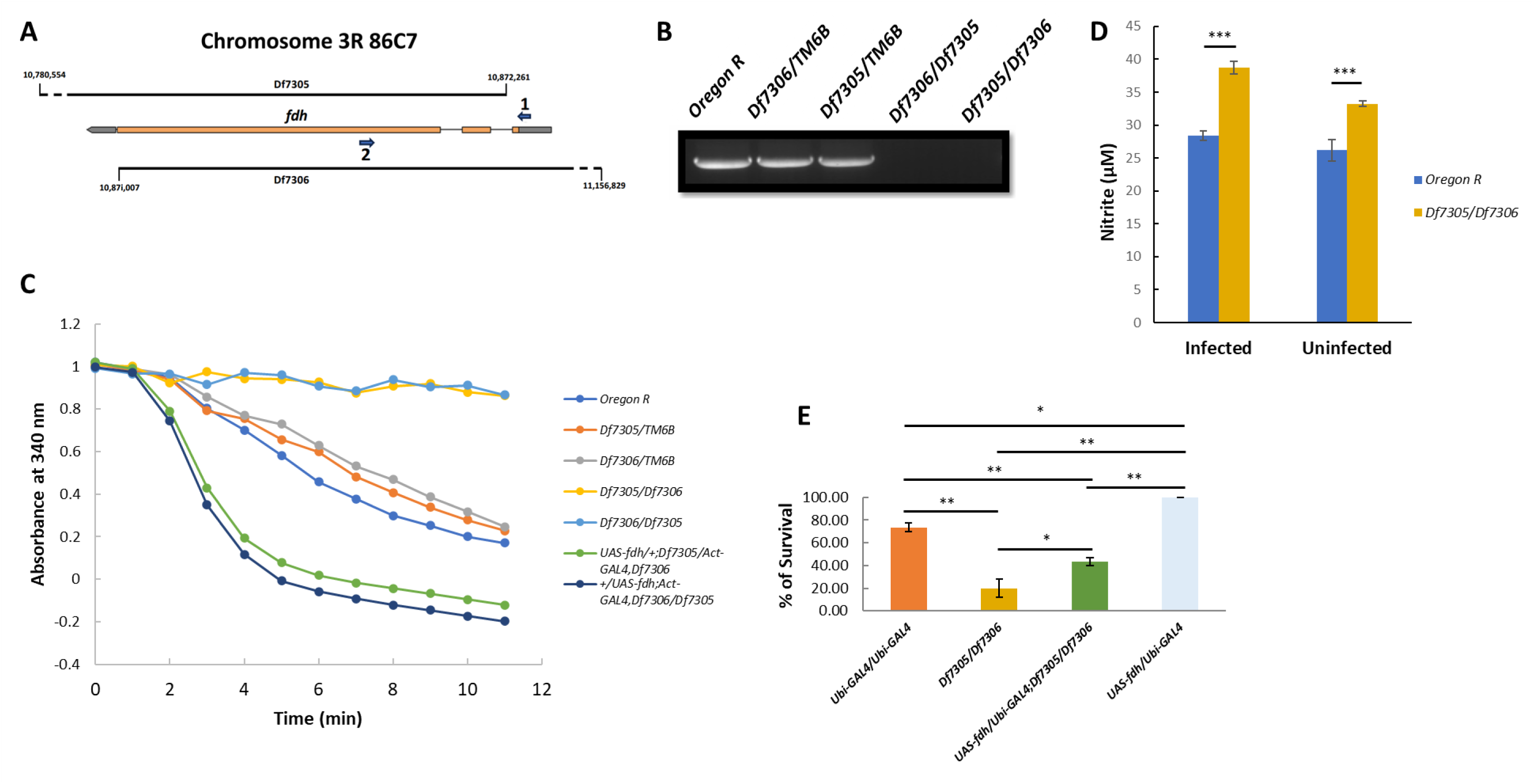
(A) Map of *fdh*. Deficiencies Df7305 and Df7306 overlap in region 86C7 of Chromosome 3 with breakpoints within the gene *fdh* that is transcribed from right to left in this diagram. The coordinates refer to bases in the *D. melanogaster* genomic sequence and are taken from FlyBase^1^. The 5’ and 3’ untranslated regions of *fdh* are shown in grey and the coding regions in orange with introns indicated by a thin line. The diagram is not drawn to scale. **(B) Flies heterozygous for Df7305 and Df7306 lack an intact copy of *fdh*.** DNA was purified from flies with the genotypes indicated and was amplified by PCR using primers 1 and 2 shown in (A). **(C) Flies heterozygous for Df7305 and Df7306 have greatly reduced Gsnor activity.** Gsnor activity in protein extracts from flies of the genotypes indicated was assayed as described in the Materials and Methods. GSNO was added at time 0 and the absorbance at 340nm was measured at ten minutes intervals thereafter. **(D) Infection by *B. bassiana* increases nitrite in flies heterozygous for Df7306 and Df7305.** The level of nitrite in extracts of wild-type (Oregon R) or heterozygous Df7305/Df7306 female flies is shown either with or without infection with *B. bassiana*. The bars show the mean ± S.E of the results of three independent experiments and the “p” value for the differences in nitrite levels between the two genotypes was calculated by one-way ANOVA and Tukey HSD tests, * indicates p ≤ 0.05 while ** indicates p ≤ 0.01. **(E) Flies with reduced Gsnor show increased sensitivity to SNP.** The SNP sensitivity of groups of 15 female flies aged for 4 to 7 days was tested as described in the Materials and Methods. The number of survivors recorded one day after exposure to SNP is shown for flies of the genotypes indicated. The data show the mean ±SE from three vials. The “p” value for the differences between the different genotypes was calculated using a one-way ANOVA (*F*3,8 = 56.68 *p*< 0.001) and a Tukey HSD test, * indicates p ≤ 0.05 while ** indicates p ≤ 0.01.

Having established these genetic tools, we challenged the *fdh* mutants with fungal pathogen *Beauveria bassiana*, the Gram-positive bacterium *Staphylococcus aureus* or the Gram-negative bacterium *Escherichia coli*. Wild type *D. melanogaster* females have been reported by others to be more sensitive to *B. bassiana* than males^24^. We have confirmed this (Fig. S2A), and the experiments reported here were performed with flies of a single sex.

Flies either heterozygous for deficiencies Df7305 and Df7306 (Df7305/Df7306) (Fig. 2A) or homozygous for the CRISPR mutation *fdh^cp2^* (Fig. S1E) are more susceptible than wild type to challenge with *B. bassiana* and *S. aureus* (Fig. 2B) and in both cases this is rescued by expression of Gsnor from an *fdh* transgene (Fig. S2A and 2B) suggesting that Gsnor is required for the activation of the Toll pathway in flies.

**Figure 2.**
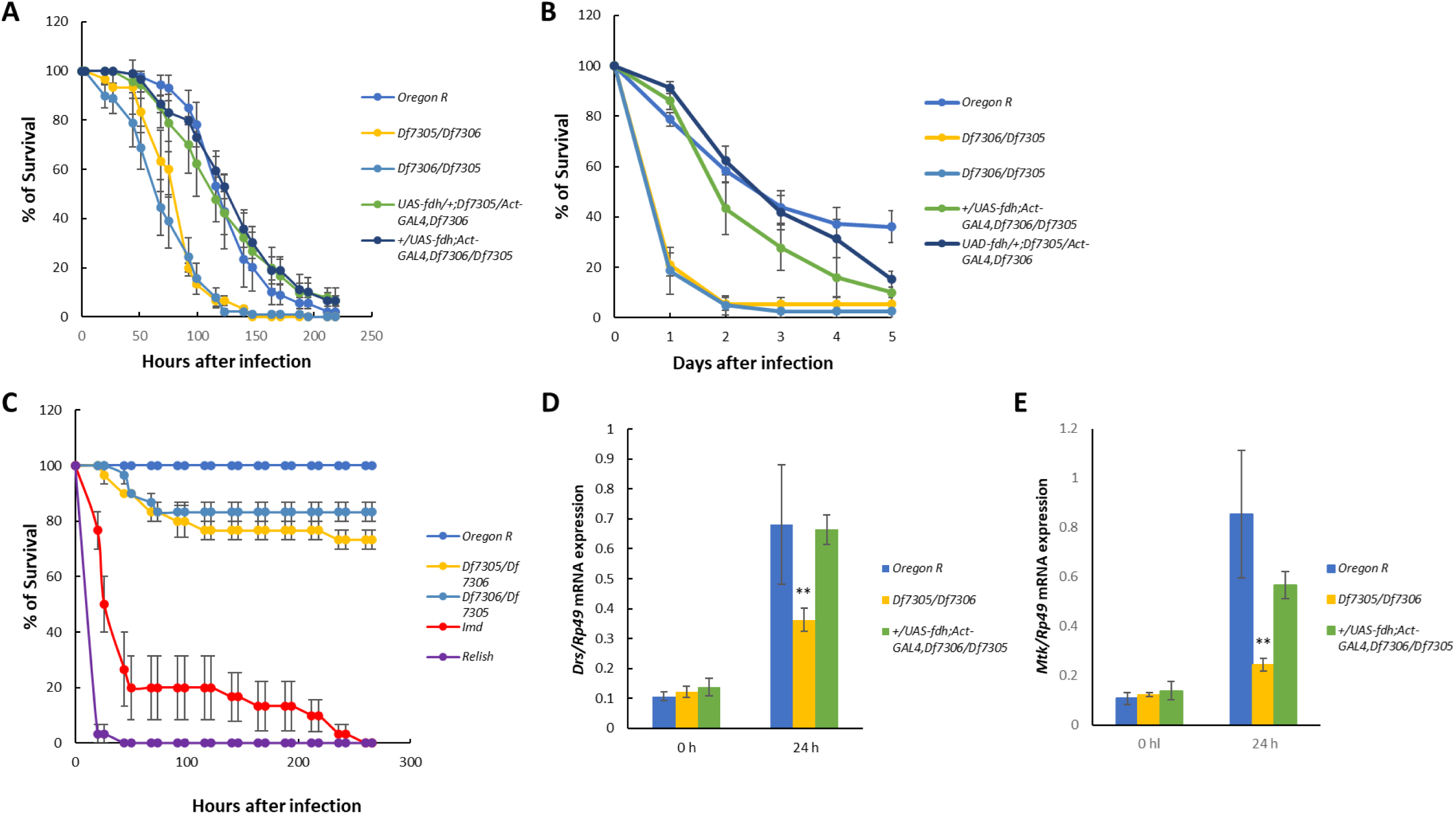
(A) Susceptibility of flies to *B. bassiana*. Groups of 15 male flies aged for 3 to 6 days, were infected with *B. bassiana* as described in the Materials and Methods. The number of flies surviving after infection was recorded every 24 hours and the survivors were transferred to a fresh vial every other day. Flies trans-heterozygous for the *fdh* deficiencies Df7305 and Df7306 were more sensitive to infection by *B. bassiana* than wild type (Oregon R). The graphs show the mean ± S.E of three independent experiments. (**B) Susceptibility of flies to infection with *S. aureus***. Groups of 30 female flies aged for 3 to 6 days were infected with a needle coated in *S. aureu*s as described in the Materials and Methods, transferred to vials containing YCMA and kept at 29°C and the number of survivors was recorded every 24 hours for five days. The graphs show the mean ± S.E of three independent experiments. **(C) Susceptibility of flies to infection with *E. coli***. About 30 female flies of the genotypes indicated were aged for 3 to 4 days then infected with *E.* coli MG1655 as derscribed in the Material and Methods. Flies homozygous for mutant alleles of *imd* (*immune deficiency*) or *rel* (*relish*) were used as controls to indicate the response of flies with defects in the IMD pathway. **(D) Activation of *Drs* and (E) *Mtk* is reduced in *fdh* mutant flies.** RNA was extracted from flies of the genotypes shown either without infection or 24 hours after infection with *B. bassiana*. The levels of *Drs* and *Mtk* relative to *Rp49* RNA were measured by Quantitative RT PCR as described in the Materials and Methods. These are shown as the mean ± S.E of the ratio of *Drs* and *Mtk* to *Rp49* RNAs from three experiments. Differences in gene expression between mutant fly lines were analysed using ANOVA followed by Fisher’s LSD (Least Significant Difference) test. Different letters indicate statistically significant differences between groups (p < 0.05).

Activation of the Toll pathway results in increased expression of the anti-microbial peptides Drosomycin (Drs) and Metchnikowin (Mtk)^3, 25, 26^ and as expected accumulation of transcripts coding for these peptides increases after infection of wild type flies with *B. bassiana* but this is reduced in *fdh* mutants (Fig. 2D and 2E).

In contrast to their response to challenge by *B. bassiana* or *S. aureus*, Df7305/Df7306 flies are no more sensitive than wild type to septic infection with the Gram-negative bacterium *E. coli* (Fig. 2C) and much less so than flies with mutations in either *IMD* or *rel* (*relish*) that code for proteins in the IMD pathway suggesting that Gsnor is not required for the activation of the IMD pathway. However, it has been suggested that NO is required for IMD activity as inhibition of nitric oxide synthesis increases the sensitivity of *D. melanogaster* larvae and adults to infection with the Gram-negative bacterium *Erwinia carotovora carotovora* ^16^.

### Genetic Epistasis Places GSNOR Function Within the Persephone “Danger-Sensing” Branch

The Toll pathway can be activated by PAMP recognition^7, 9, 27, 28, 29^ or by sensing pathogen-derived proteolytic activity via Persephone (Psh)^7, 8, 9, 10, 30^. To pinpoint where Gsnor-mediated regulation occurs within this architecture, we performed genetic epistasis analysis using mutants in the gene *necrotic* (*nec*). Nec is a serine protease inhibitor (serpin) that negatively regulates the Toll cascade, and *nec* loss-of-function mutants exhibit constitutive Toll pathway activation, resulting in spontaneous melanisation and reduced survival^10, 31, 32^. This phenotype is known to be entirely dependent on a functional Psh protease^10^.

Remarkably, we have found that the Nec phenotype is also suppressed if Gsnor activity is reduced as flies with both *nec* and *fdh* mutations show increased survival and reduced melanisation as compared with flies that are *nec* mutant but *fdh^+^*(Fig. 3A, 3B and Fig. S2D). Since Nec inhibits the cascade downstream of Psh activation, the most parsimonious model is that excessive *S*-nitrosylation directly inhibits Psh or prevents its activation, thereby blocking the signalling cascade even when the upstream serpin brake is removed.

**Figure 3.**
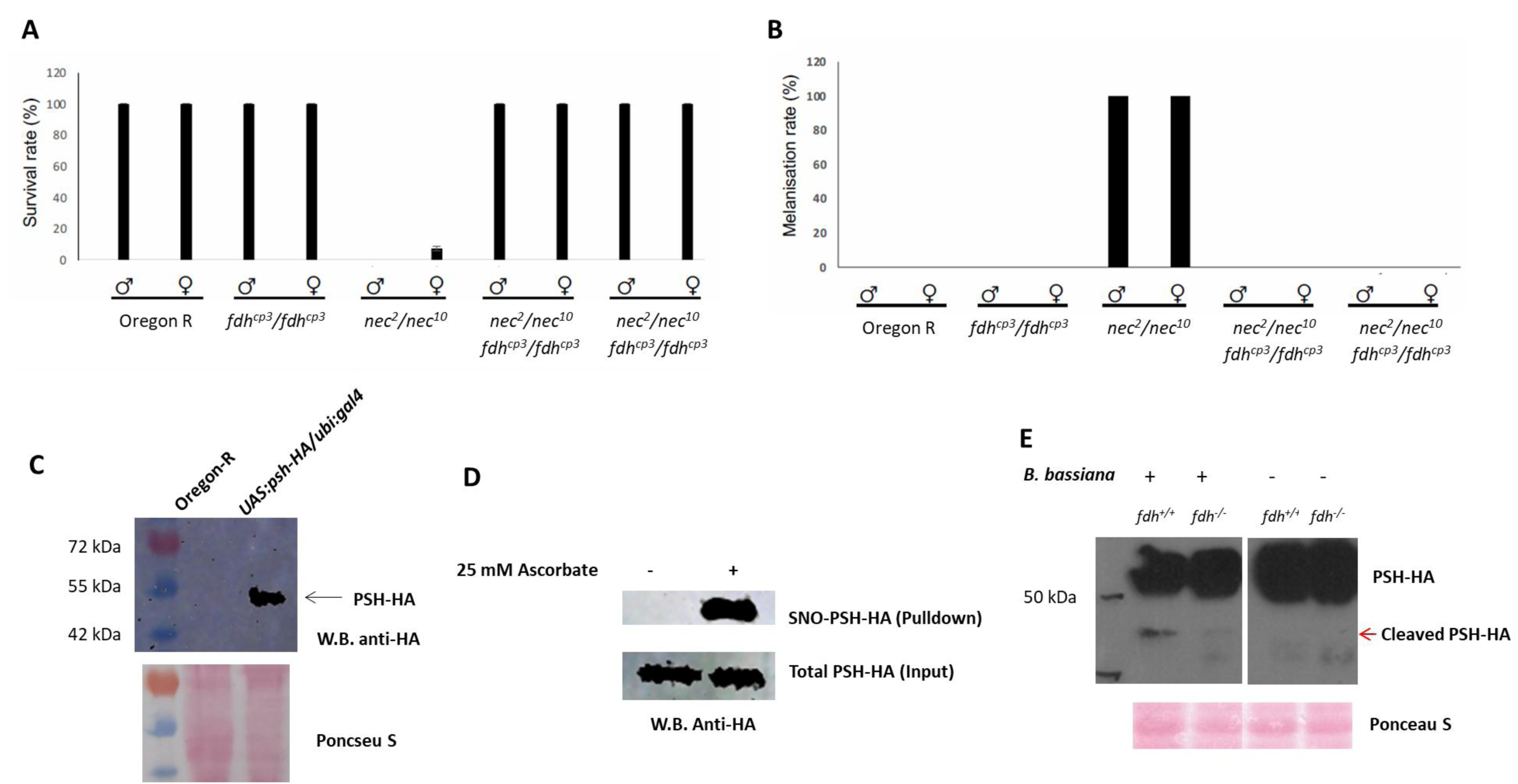
(A) Survival of *necrotic* mutant flies is suppressed by reduction in Gsnor. Total survivors were counted one day post emergence. The percentage of survivors was calculated by dividing the number of survivors by the total number of flies. The data show the mean ±SE from three repeated experiments with 24-40 flies *per* group. **(B) Melanisation of *necrotic* mutant flies is suppressed by reduction in Gsnor.** Melanised files were counted one day post emergence. The percentage of melanised flies was calculated by dividing number of melanised flies by the total number of flies. The data show the mean ±SE from three repeated experiments with 24-40 flies *per* group. **(C) Expression of HA tagged PSH in *D. melanogaster***. Total protein was extracted from wild type (Oregon R) and flies expressing Psh-HA using the UAS/GAL4 system with GAL4 expressed from the Ubiquitin promoter. Proteins were separated by SDS PAGE, transferred to a nitrocellulose membrane and HA tagged protein was detected with anti-HA antibody as described in the Materials and Methods. **(D) Persephone is S-nitrosylated *in vivo*.** Total protein extracted from 10 female flies expressing Psh-HA was submitted to the BST followed by streptavidin pulldown and Western blotting with anti-HA antibody. 25mM sodium ascorbate was used in the BST to reduce *S*-nitrosothiols and was omitted from the reaction to provide a negative control. Sample aliquots were taken before the pulldown step and used in parallel western blots to confirm that similar amounts of total protein had been loaded in each lane. **(E) Cleavage of Psh is inhibited if Gsnor is reduced.** Groups of 20 female flies, were infected by *B. bassiana* as described in the Material and Methods. Total protein was extracted from *fdh^-/-^* flies expressing Psh-HA and *fdh^+/+^* flies expression Psh-HA 3 days post *B. bassiana* inoculation. Proteins were separated by SDS PAGE, transferred to a nitrocellulose membrane and HA tagged protein was detected with anti-HA antibody as described in the Materials and Methods. Cleavage of Psh-HA (red arrow) was observed in *fdh^+/+^,* Psh-HA flies infected by *B. bassiana*, while reduced cleavage was observed in *fdh^-/-^,* Psh-HA flies infected by *B. bassiana*. No cleavage was observed in non-infected flies. Blots are cropped for clarity. Some lanes were omitted compared with the original blot; no other image processing was performed.

To further investigate whether the PAMP-recognition or danger-sensing branch was affected, we challenged flies with heat-killed pathogens. Unlike infection with live microbes, exposure to heat-inactivated *B. bassiana* (Fig. S2C) or *S. aureus* (Fig. S2B) caused little to no mortality in either wild-type or *fdh* mutant flies. This result demonstrates that the hyper-susceptibility of *fdh* mutants is not due to a defect in the recognition of structural PAMPs on the pathogen surface but is instead dependent on the presence of active virulence factors, such as secreted proteases, produced by live, metabolically active pathogens. Collectively, these genetic data strongly focussed our investigation onto the Psh-mediated danger-sensing branch of the Toll pathway as the locus of NO-mediated regulation.

### Persephone Is a Direct Target of *S*-nitrosylation *in vivo*

The genetic evidence strongly implicated the Psh protease as the key node for NO-mediated regulation. To determine if this genetic link reflects a direct biochemical interaction, we linked a human influenza haemagglutinin peptide (HA) to the C-terminus of Psh and expressed the tagged protein (Psh-HA) in *fdh^+/+^* flies using the Gal4/UAS system^21^. Expression of Psh-HA was confirmed by Western blotting with anti-HA antibody (Fig. 3C) and its *S*-nitrosylation was investigated using the Biotin Switch Technique (BST) that replaces SNO in *S*-nitrosylated proteins with biotin^33, 34^. Following application of the BST to proteins extracts of wild type flies biotinylated proteins were pulled down with streptavidin coated beads and screened for biotinylated Psh-HA in Western Blots probed with anti-HA antibody. The results show that Psh-HA can be *S*-nitrosylated *in vivo* (Fig. 3D), providing direct biochemical evidence that Psh is not merely downstream of an NO-sensitive process but is itself a *bona fide* substrate for *S*-nitrosylation within the fly. This finding forges a crucial molecular link between the whole-organism immune phenotype and the central hypothesis of direct regulation by a post-translational modification.

### *S*-nitrosylation Impedes the Proteolytic Activation of pro-Persephone

Having established that Psh can be *S*-nitrosylated, we next sought to determine the functional consequence of this modification. Previous study demonstrated *B. bassiana* effector protease PR1 cleaves pro-Psh *in vitro* between residues 115 and 116 and Psh is further processed *in vivo* by the endogenous cathepsin 26-29-p^8^. We hypothesized that *S*-nitrosylation might inhibit the Toll pathway by preventing this critical activation step. To test this, we monitored the processing of Psh-HA in wild-type (*fdh^+/+^*) and GSNOR-deficient (*fdh^cp2/cp2^*) flies following infection with *B. bassiana*.

In protein extracts from uninfected flies of either genotype, Psh-HA was detected exclusively as the full-length pro-protein. However, upon infection with *B. bassiana*, a significant portion of pro-Psh-HA in wild-type flies was cleaved, yielding a smaller, faster-migrating band corresponding to the cleaved Psh-HA. This infection-dependent processing was dramatically impaired in the *fdh^cp2/cp2^* mutants. In these flies, the amount of cleaved Psh-HA generated post-infection was markedly reduced compared to that seen in wild-type flies (Fig. 3E).

This observation was further supported by measurements of NO levels during infection. In wild-type flies, the endogenous Gsnor activity appeared sufficient to buffer any immune-induced NO production, as systemic nitrite levels remained stable after infection. In contrast, in *fdh* mutants, nitrite levels, already elevated at baseline, increased further following infection, indicating that Gsnor is essential for managing the NO surge during an immune challenge (Fig. 1D). These results provide a potential mechanistic explanation for the observed immune deficiency: pathogen infections induce NO production, the loss of Gsnor leads to an uncontrolled increase in total *S*-nitrosylation, which in turn prevents the proteolytic activation of the pro-Psh zymogen, thereby shutting down the danger-sensing arm of the Toll pathway at its point of initiation.

## Discussion

This study identifies a novel redox-based brake on the *D. melanogaster* Toll pathway, demonstrating that excessive *S*-nitrosylation prevents the proteolytic activation of the danger-sensing protease Persephone. We propose a model where Gsnor functions as a homeostatic rheostat; by metabolizing GSNO, it keeps cellular *S*-nitrosylation in check, permitting the efficient cleavage of pro-Psh required for a robust immune response to fungal and Gram-positive bacterial pathogens. In Gsnor-deficient flies, uncontrolled *S*-nitrosylation of Psh blocks this activation step, leading increased sensitivity of flies to infection by fungal and Gram-positive bacterial pathogens.

Our finding that *fdh* mutants are hyper-susceptible to the entomopathogenic fungus *B. bassiana*^35, 36, 37, 38^ provides proof-of-concept for a novel biocontrol strategy. The efficacy of mycoinsecticides is often limited by the host’s immune response^39^, and the strategy of combining microbial agents with stressors that weaken the host is a validated approach in pest management. For instance, the combination of *B. bassiana* with the insect growth regulator lufenuron, which inhibits chitin synthesis, was significantly more effective at controlling fruit fly larvae than the fungus alone^40^. We propose a similar synergistic strategy where co-application of *B. bassiana* with chemical inhibitor of Gsnor could create a potent “one-two punch”. While Gsnor is evolutionarily conserved, insect-selective Gsnor inhibitors could be deployed as adjuvants to the pathogen, simultaneously exposing insect to infection while compromising a key component of the insect’s defence, offering a rational path towards more effective and sustainable pest management solutions.

The regulation of self-amplifying serine protease cascades is fundamental for many biological processes from insect immunity to mammalian blood clotting^41, 42^. Our discovery that *S*-nitrosylation plays a key role in regulating a serine protease cascade in *D, melanogaster* may represent a conserved regulatory principle. Unlike irreversible serpin-based inhibition, regulation by a diffusible gas like NO provides a rapid, transient, and spatially localized feedback mechanism. This suggests that a “redox-protease” regulatory module may be a convergent evolutionary solution for dynamic control of proteolytic cascades, complementing traditional inhibitory mechanisms.

The versatility of NO as a signalling molecule is underscored by its opposing roles in *D. melanogaster* immunity. While our data show excessive NO is detrimental to the Toll pathway, it is required for the IMD pathway’s defence against Gram-negative bacteria¹⁶. This apparent paradox is resolved if the functional outcome is dictated by the specific molecular targets modified in each pathway. The inhibitory S-nitrosylation of pro-Psh in the Toll pathway contrasts with the presumably activating modifications of yet-unidentified targets in the IMD pathway. This allows a single molecule to differentially tune distinct arms of the immune system, suggesting NO may act as a master arbiter of immune balance, preventing immunopathology by dampening one response while another is engaged.

In conclusion, we have defined a novel regulatory axis in which the Gsnor-NO system controls Toll signalling through the direct *S*-nitrosylation of Persephone. This work not only reveals a new layer of immune homeostasis but also highlights a clear translational potential.

## ABBREVIATIONS

AMP: Anti-microbial peptide
BST: Biotein Switch Technique
DEPC: Diethyl pyrocarbonate
Drs: Drosomycin
EDTA: Ethylenediaminetetraacetic acid
*fdh*: formaldehyde dehydrogenase gene
GSNO: S*-* nitrosoglutathione
Gsnor: *S* -nitrosoglutathione reductase
HA: Haemagglutinin antigen
HEPES: (4-(2-hydroxyethyl)-1-piperazineethanesulfonic acid)
HRP: Horse radish peroxidase
Mtk: Metchnikowinu
NADH: reduced Nicotinamide adenine dinucleotide
PAMP: Pathogen Associated Molecular Patterns
PBS-T: Phosphate buffered saline + 0.1 Tween-20
PCR: Polymerase Chain Reaction
PMSF: phenylmethylsulfonyl fluoride
Psh: Persephone
*Rel*: *Relish* gene
SDS: Sodium Dodecyl Suphate
SNO: *S* -nitroso thiol
SNP: Sodium nitroprusside
*spz*: *Spätzles*
TLCK: N-alpha-tosyl-L-lysinyl-chloromethylketon
TLR: Toll like receptor
TPCK: N-tosyl-L-phenylalaninyl-chloromethylketone
Tris: Trisaminomethane
YCMA: Yeast Cornmeal Agar

## Acknowledgement

We thank Prof. Matt Tinsley (University of Stirling) for providing *Beauveria bassiana*. We thank Prof. Bruno Lemaitre (École Polytechnique Fédérale de Lausanne, Switzerland) for providing the *Relish*, *Spätzle*, and *imd* fly lines. We thank Prof. Garry Blakely (University of Edinburgh) for providing *Escherichia coli* MG1655. We are grateful to the staff of the School of Biological Sciences media and wash-up service for the supply of Drosophila media and clean glassware. This work was financially supported by the Development and Promotion of Science and Technology Talents Project (DPST) and the Institute for the Promotion of Teaching Science and Technology (IPST), Thailand, and by the Darwin Trust of the University of Edinburgh.

## Materials and Methods

### Drosophila culture

Drosophila lines were maintained at 22°C on yeast cornmeal agar medium (YCMA) (1L H_2_O, glucose 78.5g, maize meal 71.5g, Yeast 50g, agar 10.7g, Nipagen 2.7g, propionic acid 3.25ml). The genotype and origin of the strains of *D. melanogaster* used in the experiments described here are listed in Table S1. Where the flies used in an experiment are the progeny of a cross between two strains the figure indicates the genotype of the female parent first. The mutations that were not generated during this work were obtained from the sources indicated in Table S1.

### Gsnor enzyme assay

The activity of Gsnor was assayed spectrophotometrically by measuring the rate of NADH oxidation in the presence of *S*-nitrosoglutathione (GSNO). Proteins were extracted from flies in HE buffer (25 mM HEPES, 1 mM EDTA pH7.7) supplemented with the protease inhibitors [50μg/ml N-tosyl-L-phenylalaninyl-chloromethylketone (TPCK), 50μg/ml N-alpha-tosyl-L-lysinyl-chloromethylketon (TLCK) and 0.5 mM phenylmethanesulfonyl fluoride (PMSF)] and quantified using the Bradford assay (Bradford, 1976). For Gsnor activity assays 75μg of protein was incubated in 1ml of HE buffer with the addition of 350μM NADH and 350μM GSNO and NADH oxidation was measured by following the absorbance at 340nm. Zero time readings were taken immediately after the addition of GSNO.

### Measurement of nitrite in flies

Female flies were snap-frozen in liquid nitrogen and homogenised in 20μL of extraction buffer (50mM Tris-HCl pH 7.5, 150mM NaCl, 5mM EDTA, 0.1% Triton X-100) per fly using a disposable plastic grinder. Samples were centrifuged at 13,000rpm at 4°C for 15 minutes and the supernatant was transferred to a fresh pre-chilled tube. Fifty microlitres of the resulting supernatant were dispensed, in triplicate, into a flat-bottom 96-well enzymatic assay plate, after which 50 μL of Sulfanilamide Solution from the Griess Reagent System (Promega, G2930) were added. Plates were incubated at room temperature for 10 min, protected from light, before the sequential addition of 50μL N-1-naphthylethylenediamine dihydrochloride Solution. After a further 10 min incubation in the dark, the azo chromophore was quantified at 540 nm using a microplate reader within 30 min of colour development. Sodium nitrite standards (0–100 μM, prepared in homogenate buffer) were processed in parallel, and sample nitrite concentrations were interpolated from the standard curve generated on the same plate.

### Sodium nitroprusside (SNP) sensitivity assay

Female flies aged between 3 and 6 days were transferred to empty vials and starved for two hours at 25°C and then transferred to vials containing a cotton roll soaked in 5% sucrose solution supplemented with 5 mM SNP. The vials were kept at 25°C and the number of survivors was recorded one day after the treatment.

### Septic infection with bacteria

*Staphylococcus aureus* NCTC 8325 (https://www.culturecollections.org.uk) was used to test sensitivity to Gram positive bacteria and *Escherichia coli* strain MG1655^1^ was used to test sensitivity of Gram negative bacteria. Bacteria were grown on LB (10g/L Foremedium Tryptone, 5g/L Foremedium Yeast extract, 10g/L NaCl) agar (15g/L) plates and for each experiment a single colony was transferred to 5ml LB and grown overnight at 37°C shaking at 200rpm and then the culture was centrifuged. Flies were anaesthetised with CO_2_ or by chilling on ice and their abdomen was pierced with a tungsten needle that had been dipped in the bacterial pellet. After infection the flies were transferred to vials containing YCMA and kept at 29°C. The proportion of flies surviving was calculated at the time points shown in the relevant figures with any flies that died within two hours excluded from the total number at the start of the experiment.

### Assaying sensitivity to infection with *B. bassiana*

*Beauveria bassiana* was grown at 29°C on potato dextrose agar (39g PDA (Sigma P2182) in 1L of water) supplemented with 50μg/ml chloramphenicol. Once the mycelium had covered the plate spore formation was induced by protecting the plates from light and leaving them in a fume hood until completely desiccated. The dried plates were then stored in the dark at 4°C. For infection assays twenty 3-7 days old flies were anaesthetized by chilling on ice and transferred to a 2ml microcentrifuge tube containing *B. bassiana* with mycelium and spores scraped from an area of about 1cm^2^ of a dried plate. The tubes were then shaken gently by hand for 2 minutes after which the flies were transferred to vials containing YCMA, maintained at 29°C and the number of surviving flies was recorded over time. Flies that died within two hours of infection were excluded when calculating survival rates.

### Genomic DNA extraction from Drosophila

Thirty flies were frozen at −70°C and then homogenised with a disposable plastic grinder in 400μL of 100mM Tris-HCl pH7.5, 100mM EDTA, 100mM NaCl, 0.5% SDS (Sodium Dodecyl Sulphate). The homogenate was incubated at 65°C for 30 minutes and then 800μL of 1.67M C_2_H_3_O_2_K, 4M LiCl were added to the tubes, followed by incubation on ice for 10 minutes and a subsequent centrifugation at 14,000rpm for 15 minutes. Equal volumes of the supernatant were transferred to two clean tubes, and DNA was precipitated by adding 700μL isopropanol followed by centrifugation at 14,000rpm for 15 minutes. The pellet was then dissolved in 100μL sterile distilled water.

### Detection of *fdh* sequences by PCR

Genomic DNA of flies heterozygous for the overlapping deletions Df7305 and Df7306 was amplified by PCR to confirm that they do not contain an intact *fdh*. The reaction mix contained 1 x *Ex Taq* Buffer (TakaraBio), 0.2mM dNTPs, 0.2µM of each of primer 1 (5‘-AATAAACCATACTGCAAAGATGTCTGCTAC-3’ and primer 2 (5‘-TGCAGCTGAGACGG-3’) (Fig. 1A), 5%v/v genomic DNA, and 0.05unit/µL TaKaRa Ex Taq DNA polymerase. The DNA was denatured at 94°C for 1 minute followed by 5 cycles at 94°C for 0.5 minute, 60°C for 0.5 minute and 72°C for 2.5 minutes, and then 25 cycles of 94°C for 0.5 minute and 68°C for 2.5 minutes. Amplified DNA was analysed by agarose gel electrophoresis (1% agarose in 1 x Tris-acetate-EDTA buffer [40 mM Tris, 20 mM Acetic acid and 1 mM EDTA]). Gels were stained with 0.5μg/ml ethidium bromide and the DNA visualized using an ultraviolet imaging system

### Total RNA extraction

Five adult female flies were frozen in liquid nitrogen and then homogenised in 500μL of TRizol (Invitrogen) using a disposable plastic grinder. The homogenate was left at room temperature for 5 minutes followed by centrifugation at 12,000rpm for 10 minutes at 4°C. 180μL of the supernatant were transferred to a new microcentrifuge tube and 60μL of chloroform was added. The homogenate and chloroform were mixed vigorously by hand followed by incubation at room temperature for 3 minutes and centrifugation at 10,000rpm for 15 minutes at 4°C. About 80μL of the upper phase was transferred to a new microcentrifuge tube and 100μL of isopropanol was added followed by incubation at room temperature for 5 minutes and centrifugation at 12,000rpm for 10 minutes at 4°C. The supernatant was replaced by 600μL of 75% ethanol followed by centrifugation at 2,200g for 5 minutes at 4°C. After removing the supernatant, the RNA pellet was dried in a laminar flow cabinet and then dissolved in 55μL of diethylpyrocarbonate (DEPC) treated water.

### RNA extraction and RT-PCR

Twenty five 3-4 days old flies were frozen in liquid nitrogen and subsequently ground in 500µL of TRizol (Invitrogen) using a bead mill (Qiagen). The homogenate was left at room temperature for 5 minutes before centrifugation at 5,600g for 10 minutes at 4°C. The supernatant was transferred to a new microcentrifuge tube and 100µL of chloroform was added. The contents were mixed vigorously by hand and incubated at room temperature for 3 minutes before centrifugation at 10,000g for 15 minutes at 4°C. The upper phase was transferred to a new microcentrifuge tube and 250µL of isopropanol was added followed by incubation at room temperature for 10 minutes and centrifugation at 12,000g for 10 minutes at 4°C. The supernatant was replaced with 0.5mL of 75 % ethanol followed by centrifugation at 7,500xg for 5 minutes at 4°C. The supernatant was removed, and the RNA pellet was dried in a laminar flow cabinet and then dissolved in 50µL of diethylpyrocarbonate (DEPC) treated water.

### Quantitative RT-PCR

Expression of the anti-microbial peptide encoding genes *Drosomycin* (*Drs*) and *Metchnikowin* (*Mtk*) was quantified by reverse transcript polymerase chain reaction (RT PCR) using SYBR Green I Master Mix (Thermo Fisher Scientific) in a LightCycler 480 system (Roche). Relative gene expression levels were quantified using the 2^-ΔΔCt^ method^2^ the housekeeping gene *Rp49* (Ribosomal protein 49) serving as an internal control for normalisation. The primers and conditions used to quantify the expression of these genes have been described previously^3,4^ The primers used were 5’-AGATCGTGAAGAAGCGCACCAAG-3’and 5’-CACCAGGAACTTCTTGAATCCGG-3’for *Rp49*, 5‘-CGTGAGAACCTTTTCCAATTATGATG-3’and 5’-TGGTGGAGTTGGGCTTCATG-3’for *Drs*, and 5-GATGCAACTTAATCTTGGAGCG-3’and 5’-TTAATAAATTGGACCCGGTCTTGGTTGG-3’for *Mtk*.

### Total protein extraction and quantification

Flies were homogenised in 20μL of extraction buffer (50mM Tris-HCl pH 7.5, 150mM NaCl, 5mM EDTA, 0.1% Triton X-100) *per* fly freshly supplemented with protease inhibitors (50μg/ml TPCK 50 μg/ml TLCK and 0.5mM PMSF). Samples were centrifuged at 13,000rpm at 4°C for 15 minutes and the supernatant was transferred to a fresh pre-chilled tube. Protein concentrations were measured using the Bradford Assay^5^.

### SDS-Poly-Acrylamide Gel Electrophoresis (SDS-PAGE)

Protein samples were mixed with a 4x stock of sample buffer to a final concentration of 50mM Tris-HCl pH 6.8, 2% SDS, 0.02% bromophenol blue and 10% glycerol either with or without 50mM dithiothreitol. Samples were heated at 85°C for 10 min before separation by polyacrylamide gel electrophoresis. Gels were washed in H_2_O before being incubated in staining solution (0.25% Coomassie Brilliant Blue R, 40% methanol, 7% acetic acid) for between 30 min and one hour. Gels were de-stained overnight in de-staining solution (40% methanol, 10% acetic acid) and then photographed

### Western Blots

Proteins were transferred to nitrocellulose membranes using a Bio-Rad Trans Blot® system either overnight at a constant voltage of 20V or for 2-3 hours at 90V. Proteins were visualised on the membranes with Ponceau S (0.1% Ponceau S, 5% acetic acid) for 1 min and then rinsed with H_2_O before being photographed. Membranes were de-stained with PBS-T (137mM NaCl, 2.7mM KCl, 10mM Na_2_HPO_4_, 1.8mM KH_2_PO_4_, 0.1% Tween-20) and blocked for 1 hour at room temperature with 5% dried skimmed milk in PBS-T before incubation with primary antibodies either overnight at 4°C or at room temperature for 1-2 hours. After washing to remove excess primary antibody the membrane was incubated for 1 hour at room temperature with the appropriate secondary antibody coupled to horseradish peroxidase (HRP). SuperSignal West Pico/Dura Chemiluminescent Substrate (Thermo Scientific) was added to the membranes and labelled proteins were detected with X-ray film. All antibodies were diluted in 5% dried skimmed milk in PBS-T. The primary antibodies were mouse anti-HA (Roche, clone 12CA5) and HRP conjugated goat anti-Biotin (Cell Signalling). The secondary antibody for detection of HA antigen was HRP linked goat anti-mouse IgG HRP (Cell Signalling #7075).

### Expression of Gsnor from *fdh* transgenes and expressing Psh-HA

Gsnor was expressed in flies using the UAS/GAL4 system^6^ to drive expression of an *fdh* transgene linked to the GAL4 UAS binding site. GAL4 protein was expressed from either the *ubi* (ubiquitin) or *act5C* (Actin 5C) promoter.

The coding sequence of *psh* was amplified using Phusion high-fidelity polymerase (New England Biolabs, UK) from freshly synthetized cDNA of wild type Oregon- R flies. The primers (Forward: 5’-CACCATGCCATTGAAGTGGTC-3’ and Reverse: 5’-TTACTTCACCCGATTGTCCGG-3’) were designed to add the nucleotides CACC at the 5’end of the coding strand to allow TOPO® cloning (Life Technologies). The PCR products were gel-purified and cloned into the pENTRTM/D-TOPO® vector according to the manufacturers’ instructions, transfected into *E.coli* and plated on LB agar containing 50µg/ml kanamycin.

Plasmid DNA was isolated from single colonies and sequenced. Inserts from positive constructs were transferred by Gateway® cloning (LR reactions following manufacturer’s instructions - Life Technologies) into pUASt-HA to generate pUASt-*Psh*-HA^7^ for expression of C-terminal HA-tagged protein in *D. melanogaster*. Recombinant clones were selected on LB agar containing 50µg/ml ampicillin and were confirmed by sequencing. pUASt-*Psh*-HA constructs were purified using the QIAfilter plasmid midi kit (Qiagen), in accordance with the manufacturer’s instructions. DNA quality and concentrations were measured using a NanoDrop spectrophotometer ND 1000, and 50μg of each construct was sent to Genetic Services Inc for integration at the attP40 φC31 integration site on the 2^nd^ chromosome^8^.

### Detection of *S-*nitrosylated PSH-HA

The Biotin Switch Technique (BST)^9,10^ was used to detect *S*-nitrosylation of Psh-HA. Flies expressing Psh-HA were homogenised in extraction buffer (100mM HEPES pH7.8, 1mM EDTA, 0.1mM Neocuproine, 0.5% Triton X-100, 50μg/ml TPCK, 50μg/ml TLCK and 0.5 mM PMSF. Samples were then centrifuged at 13,000rpm at 4°C for 15 minutes. Free thiols on proteins in the supernatant were blocked in an equal volume of blocking buffer (250mM HEPES pH7.8, 1mM EDTA, 0.1 mM Neocuproine, 5% (w/v) SDS, 50mM N-Ethylmaleimide) for 30 minutes at 55°C. The blocking buffer was removed by precipitating proteins with two volumes of cold acetone. After 20 minutes at −20°C, samples were centrifuged at 15,000rpm for five minutes at 4°C. The pellet was washed three times with 70% acetone and resuspended in 85μL of 250mM HEPES pH7.8, 1 mM EDTA, 0.1 mM Neocuproine, 1% (w/v) SDS). *S*-nitrosothiols were reduced by adding sodium ascorbate to a final concentration of 25mM and the newly free thiols were biotinylated by adding biotin-*N*-[6-(biotinamido)hexyl]-3′-(2′-pyridyldithio)-propionamide) to a final concentration of 0.4mM placed on a rocker plate for one hour. Ascorbate was omitted from samples used as negative controls. Proteins were then precipitated and washed with acetone as described above.

For detection of biotinylated Psh-HA the pellet was dissolved in 300μL of 25mM HEPES, pH7.8, 1mM EDTA, 0.1mM Neocuproine, 1% (w/v) SDS to which 20µL of streptavidin beads in 100µL of 25 mM HEPES, 1mM EDTA, 0.1mM Neocuproine, 100mM NaCl, 0.5% Triton X-100, were added and incubated at 4°C on a rocking plate overnight. Next morning samples were washed five times with 500µL of 25mM HEPES, 1 mM EDTA, 0.1 mM Neocuproine, 600mM NaCl, 0.5% Triton X-100 and resuspended in 20µL of elution buffer (25mM HEPES, 1mM EDTA, 0.1mM Neocuproine, 1% v/v β-mercaptoethanol). After 30 minutes at room temperature the beads were pelleted by centrifugation at maximum speed for one minute at room temperature and 30µl of the supernatant was loaded on a SDS-PAGE and biotinylated Psh-HA was detected in a Western blot using anti-HA antibody.

### CRISPR/Cas9 deletion of *fdh*

The *fdh* gene was replaced with DNA coding for the visible marker DsRed using CRISPR/Cas9 to cleave DNA on either side of *fdh* and a plasmid with *DsRed* flanked by sequences from either side of *fdh* as the template to allow homology-directed repair (Fig S1A). The “CRISPR Optimal Target Finder” web tool (https://flycrispr.org/target-finder/)^11^ was used to find targets for CRISPR/Cas9 cleavage on either side of *fdh*. The gRNA target sequences identified were Chromosome 3R nucleotides 10870883 to 10870905 to the left of *fdh* and nucleotides 10872425 to 10872446 to the right of *fdh.* Two homology arms adjacent to the gRNA target sequences and flanking *fdh* were amplified by PCR using primers with restriction sites allowing the products to be inserted on either side of the 3x*DsRed* coding DNA in the plasmid pHD-DsRed-attP-w^+^ (Addgene #80898 donated by K. O’Connor-Giles) to give the plasmid pHD-DsRed-attP-*fdh*. The primers, with restriction sites underlined, for amplifying the sequence to the left of *fdh* were 5’-CA<u>CTGCAG</u>CGTATCTCTACGGATATCC-3’ and 5’- CA<u>AGATCT</u>TTGGGGGTCGGATTACTGTC-3’, and for amplifying the sequence on the right of *fdh* were 5’-ATA<u>CATATG</u>AAGCTCACCGGGACTCAG-3’and 5’-GATA<u>GAATTC</u>ACACTGACGATGTGATCCACATAG-3’.

A plasmid expressing gRNAs to direct cleavage to the left and right of *fdh* was constructed from pCFD4-U6:1_U6:3tandemgRNAs (Addgene #49411 donated by S. Bullock) as described by Port et al^12^. Primers containing gRNA sequences for cleavage on either side of *fdh* were used to amplify a fragment that was then cloned into BbsI cut pCFD4 by homology directed cloning (Gibson Assembly® Cloning Kit – New England Biolabs. Primers for this PCR amplification, with the gRNA sequence underlined, were 5’-TATATAGGAAAGATATCCGGGTGAACTTCG<u>ACATAAGAGTATCTTCAT</u>TGGTTTTAGAGCTA GAAATAGCAAG-3’, and 5’-ATTTTAACTTGCTATTTCTAGCTCTAAAAC<u>TAAAAGCTCACCGGGACTC</u>CGACGTTAAATTGA AAATAGGTC-3’). The resulting plasmid pCFD4-gRNAs expresses one gRNA from the U6:1 promoter and the other from the U6:3 promoter.

Both constructs pCFD4-gRNAs and pHD-DsRed-attP-*fdh* were purified and sent to the Genetic services Inc, where they were co-injected into embryos of *D. melanogaster* strain BDSC #51323 (*y^[1]^,*{*vas-Cas9*}ZH*-*2A *w*^1118^/FM7) that expresses Cas9 protein under the control of the *vasa* promoter. Transgenic flies expressing DsRed were crossed with a balancer strain to construct stocks homozygous for the *fdh* deletion and to remove the chromosomes carrying *vas-Cas9*. The replacement of *fdh* by *DsRed* was confirmed by PCR and sequencing of genomic DNA using primers fwd - 5’-CTTGGAGCCGTACTGGAACTG 3’ and rev - 5‘-GCCTCCGATTTGGTTTGTTG-3‘as shown in Fig S1B.

---

**Figure S1.**
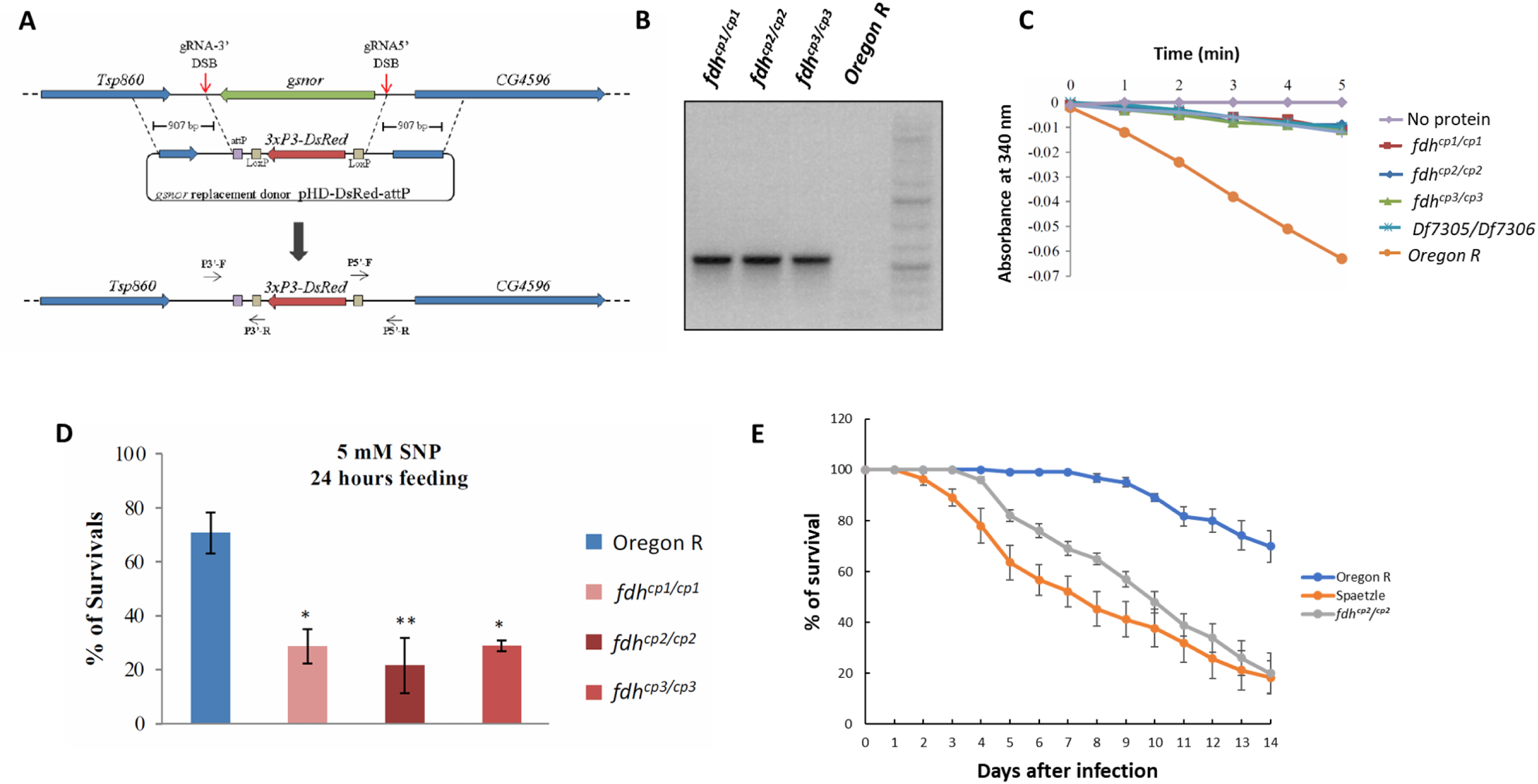
(A) Strategy for replacing *fdh* coding sequence with *DsRed*. The method used is described in Materials and Methods. The chromosomal region containing *fdh* gene and flanking genes is diagrammed in the top line. gRNAs expressed from plasmid pCFD4-gRNA direct Cas9 nuclease to cut intergenic DNA on either side of *fdh*. The resulting gap is repaired by the homology directed repair pathway using the pHD-DsRed-attP plasmid as the homology template thereby replacing *fdh* with 3xP3-*DsRed*. This diagram is not drawn to scale. **(B) Confirmation that *DsRed* is integrated in place of *fdh*.** PCR using genomic DNA from flies homozygous for mutations *fdh^cp1^, fdh^cp2^*, *fdh*^cp3^ and Oregon R flies as template and the primers P5’-F and P5’-R (Fig. S1A) confirmed the integration of *DsRed* in the mutant alleles. **(C) Flies homozygous for CRISPR replacement of *fdh* have reduced Gsnor activity.**Protein extracts from flies homozygous for *fdh^cp1^, fdh^cp2^* or *fdh^cp3^* were assayed for Gsnor activity as described in the Materials and Methods. Protein from flies heterozygous for Df7305 and Df7306 was included for comparison, with protein from Oregon R flies used as a positive control and a reaction without protein as negative control. **(D) Flies homozygous for CRISPR generated mutations of *fdh* show increased sensitivity to SNP.** The SNP sensitivity of groups of 20 female flies aged for 3 to 6 days was tested as described in the Materials and Methods. The data represent the mean ±SE of three independent experiments and the results were analysed by one-way ANOVA and Tukey HSD tests, * indicates p<0.05 and ** indicates p<0.01. **(E). Flies homozygous for CRISPR generated mutations of *fdh* show increased susceptibility to *B. bassiana***. Groups of 15-20 flies (3-7 days old) of the indicated genotypes were infected with *B. bassiana* as described in Materials and Methods. Each experiment used three vials of 15-20 flies each (45-60 flies total per genotype per experiment). Survival was monitored daily for 14 days post-infection. The survival curves show pooled data from three independent experiments (total n = 135-180 flies per genotype).

**Figure S2.**
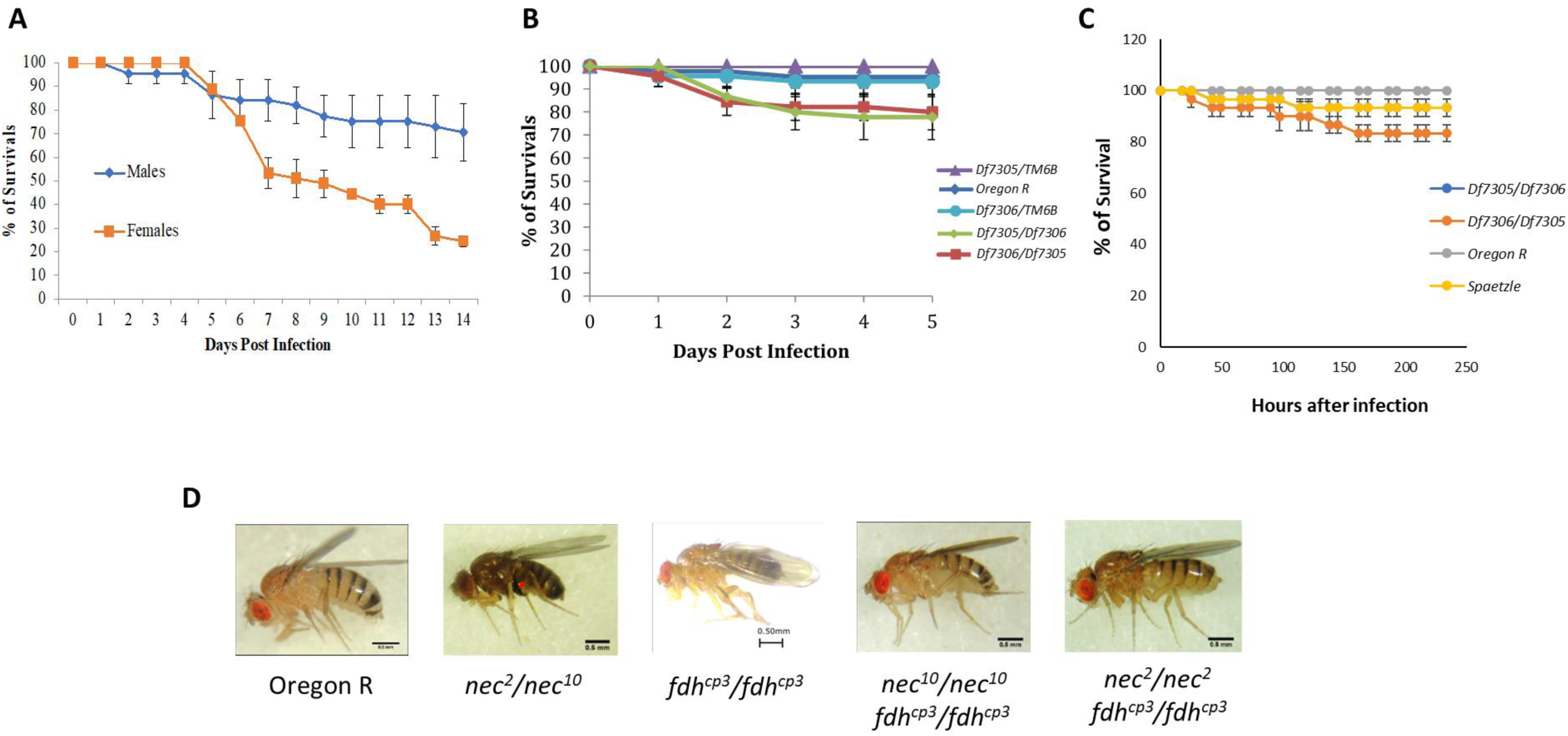
(A) Female flies are more susceptible than males to infection with *B. bassiana*. Groups of 15 male or female wild type (Oregon R) flies, aged for 3 to 6 days, were tested for their sensitivity to *B. bassiana* infection as described in the Materials and Methods. The figure shows the mean ±SE of three repetitions. **(B) Flies with reduced Gsnor are not affected by exposure to heat killed *S. aureus*.** Groups of 15 female flies, aged for 3 to 6 days were pierced with a needle coated in heat-killed *S. aureus*, placed in vials of fresh YCMA, incubated at 29°C and the number of survivors was recorded every 24 hours for five days. The figure shows the mean ±SE of three independent experiments. **(C) Flies with reduced Gsnor are not affected by exposure to heat killed *B. bassiana*.** Flies were exposed to heat killed spores of *B. bassiana* as described for live spores in the Materials and Methods and the percentage of surviving flies recorded over time. The figure shows the mean ±SE of three independent experiments. **(D) Melanisation of flies due to *nec* mutations is suppressed in flies with reduced Gsnor activity.** Each panel shows a 3 to 5 days old female fly of the genotypes indicated.

**Table S1:**
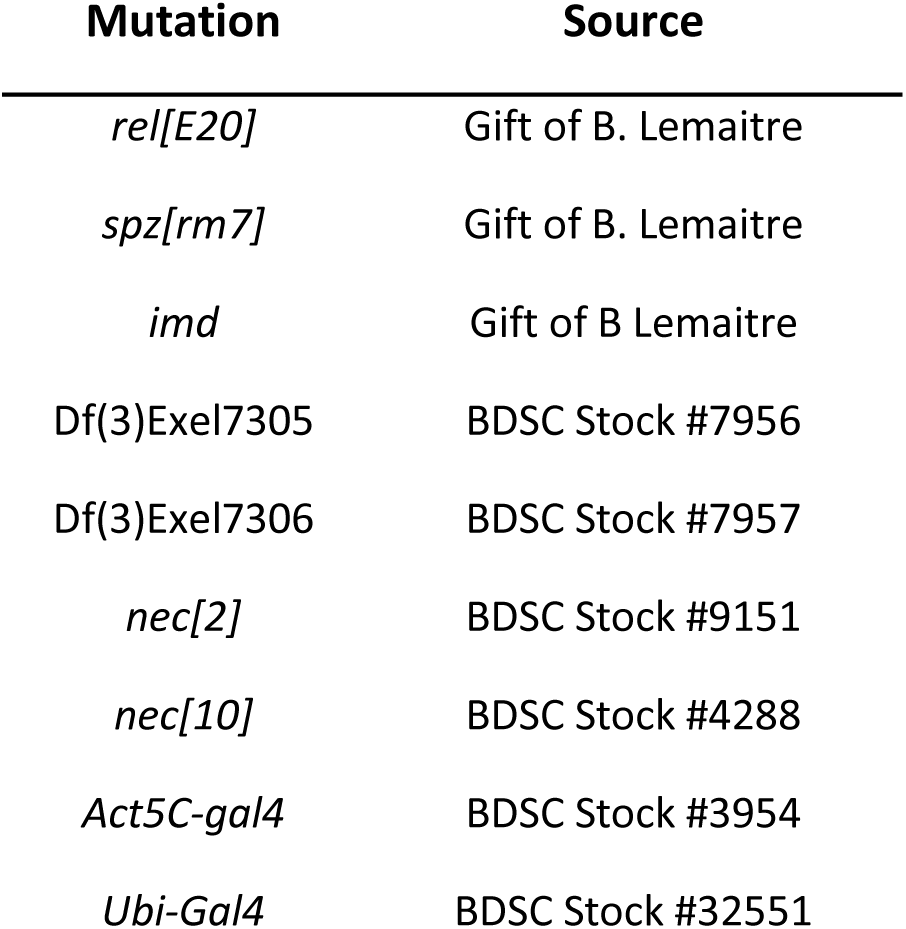
The sources of mutant alleles used in these experiments.ss.

## Reference

1. Bernheim A, Cury J, Poirier EZ. The immune modules conserved across the tree of life: Towards a definition of ancestral immunity. PLoS Biol 22, e3002717 (2024).

2. Buchon N, Silverman N, Cherry S. Immunity in Drosophila melanogaster--from microbial recognition to whole-organism physiology. Nat Rev Immunol 14, 796–810 (2014).

3. Lemaitre B, Nicolas E, Michaut L, Reichhart JM, Hoffmann JA. The dorsoventral regulatory gene cassette spätzle/Toll/cactus controls the potent antifungal response in Drosophila adults. Cell 86, 973–983 (1996).

4. Lemaitre B, et al. A recessive mutation, immune deficiency (imd), defines two distinct control pathways in the Drosophila host defense. Proc Natl Acad Sci U S A 92, 9465–9469 (1995).

5. Lemaitre B, Hoffmann J. The host defense of Drosophila melanogaster. Annu Rev Immunol 25, 697–743 (2007).

6. Hoffmann JA, Reichhart JM. Drosophila innate immunity: an evolutionary perspective. Nat Immunol 3, 121–126 (2002).

7. Gottar M, et al. Dual detection of fungal infections in Drosophila via recognition of glucans and sensing of virulence factors. Cell 127, 1425–1437 (2006).

8. Issa N, et al. The Circulating Protease Persephone Is an Immune Sensor for Microbial Proteolytic Activities Upstream of the Drosophila Toll Pathway. Mol Cell 69, 539–550.e536 (2018).

9. El Chamy L, Leclerc V, Caldelari I, Reichhart JM. Sensing of ‘danger signals’ and pathogen-associated molecular patterns defines binary signaling pathways ‘upstream’ of Toll. Nat Immunol 9, 1165–1170 (2008).

10. Ligoxygakis P, Pelte N, Hoffmann JA, Reichhart JM. Activation of Drosophila Toll during fungal infection by a blood serine protease. Science 297, 114–116 (2002).

11. Coleman JW. Nitric oxide in immunity and inflammation. Int Immunopharmacol 1, 1397–1406 (2001).

12. Lundberg JO, Weitzberg E. Nitric oxide signaling in health and disease. Cell 185, 2853–2878 (2022).

13. Hess DT, Matsumoto A, Kim SO, Marshall HE, Stamler JS. Protein S-nitrosylation: purview and parameters. Nat Rev Mol Cell Biol 6, 150–166 (2005).

14. Feechan A, Kwon E, Yun BW, Wang Y, Pallas JA, Loake GJ. A central role for S-nitrosothiols in plant disease resistance. Proc Natl Acad Sci U S A 102, 8054–8059 (2005).

15. Liu L, Hausladen A, Zeng M, Que L, Heitman J, Stamler JS. A metabolic enzyme for S-nitrosothiol conserved from bacteria to humans. Nature 410, 490–494 (2001).

16. Foley E, O’Farrell PH. Nitric oxide contributes to induction of innate immune responses to gram-negative bacteria in Drosophila. Genes Dev 17, 115–125 (2003).

17. Yun BW, et al. S-nitrosylation of NADPH oxidase regulates cell death in plant immunity. Nature 478, 264–268 (2011).

18. Hou Q, et al. Nitric oxide metabolism controlled by formaldehyde dehydrogenase (fdh, homolog of mammalian GSNOR) plays a crucial role in visual pattern memory in Drosophila. Nitric Oxide 24, 17–24 (2011).

19. Dijkers PF, O’Farrell PH. Dissection of a hypoxia-induced, nitric oxide-mediated signaling cascade. Mol Biol Cell 20, 4083–4090 (2009).

20. Parks AL, et al. Systematic generation of high-resolution deletion coverage of the Drosophila melanogaster genome. Nat Genet 36, 288–292 (2004).

21. Brand AH, Perrimon N. Targeted gene expression as a means of altering cell fates and generating dominant phenotypes. Development 118, 401–415 (1993).

22. Gratz SJ, Rubinstein CD, Harrison MM, Wildonger J, O’Connor-Giles KM. CRISPR-Cas9 Genome Editing in Drosophila. Curr Protoc Mol Biol 111, 31.32.31–31.32.20 (2015).

23. Pavlopoulos A, Averof M. Establishing genetic transformation for comparative developmental studies in the crustacean Parhyale hawaiensis. Proc Natl Acad Sci U S A 102, 7888–7893 (2005).

24. Taylor K, Kimbrell DA. Host immune response and differential survival of the sexes in Drosophila. Fly (Austin) 1, 197–204 (2007).

25. Lemaitre B, Reichhart JM, Hoffmann JA. Drosophila host defense: differential induction of antimicrobial peptide genes after infection by various classes of microorganisms. Proc Natl Acad Sci U S A 94, 14614–14619 (1997).

26. Levashina EA, Ohresser S, Bulet P, Reichhart JM, Hetru C, Hoffmann JA. Metchnikowin, a novel immune-inducible proline-rich peptide from Drosophila with antibacterial and antifungal properties. Eur J Biochem 233, 694–700 (1995).

27. Buchon N, et al. A single modular serine protease integrates signals from pattern-recognition receptors upstream of the Drosophila Toll pathway. Proc Natl Acad Sci U S A 106, 12442–12447 (2009).

28. Pili-Floury S, et al. In vivo RNA interference analysis reveals an unexpected role for GNBP1 in the defense against Gram-positive bacterial infection in Drosophila adults. J Biol Chem 279, 12848–12853 (2004).

29. Gobert V, et al. Dual activation of the Drosophila toll pathway by two pattern recognition receptors. Science 302, 2126–2130 (2003).

30. Dudzic JP, Hanson MA, Iatsenko I, Kondo S, Lemaitre B. More Than Black or White: Melanization and Toll Share Regulatory Serine Proteases in Drosophila. Cell Rep 27, 1050–1061.e1053 (2019).

31. Green C, Levashina E, McKimmie C, Dafforn T, Reichhart JM, Gubb D. The necrotic gene in Drosophila corresponds to one of a cluster of three serpin transcripts mapping at 43A1.2. Genetics 156, 1117–1127 (2000).

32. Levashina EA, et al. Constitutive activation of toll-mediated antifungal defense in serpin-deficient Drosophila. Science 285, 1917–1919 (1999).

33. Jaffrey SR, Snyder SH. The biotin switch method for the detection of S-nitrosylated proteins. Sci STKE 2001, pl1 (2001).

34. Forrester MT, Foster MW, Benhar M, Stamler JS. Detection of protein S-nitrosylation with the biotin-switch technique. Free Radic Biol Med 46, 119–126 (2009).

35. Dong Y, Morton JC, Jr., Ramirez JL, Souza-Neto JA, Dimopoulos G. The entomopathogenic fungus Beauveria bassiana activate toll and JAK-STAT pathway-controlled effector genes and anti-dengue activity in Aedes aegypti. Insect Biochem Mol Biol 42, 126-132 (2012).

36. Lee JY, Woo RM, Choi CJ, Shin TY, Gwak WS, Woo SD. Beauveria bassiana for the simultaneous control of Aedes albopictus and Culex pipiens mosquito adults shows high conidia persistence and productivity. AMB Express 9, 206 (2019).

37. Al Khoury C, Nemer N, Bernigaud C, Fischer K, Guillot J. First evidence of the activity of an entomopathogenic fungus against the eggs of Sarcoptes scabiei. Vet Parasitol 298, 109553 (2021).

38. Darbro JM, Graham RI, Kay BH, Ryan PA, Thomas MB. Evaluation of entomopathogenic fungi as potential biological control agents of the dengue mosquito, Aedes aegypti (Diptera: Culicidae). Biocontrol Science and Technology 21, 1027–1047 (2011).

39. Wang Y, Cui C, Wang G, Li Y, Wang S. Insects defend against fungal infection by employing microRNAs to silence virulence-related genes. Proc Natl Acad Sci U S A 118, (2021).

40. Ndii A, Rahardjo B, Himawan T. The Combination of Entomopathogenic Fungus of Beauveria bassiana (Balls) Vuill. with the Insect Growth Regulator (IGR) of Lufenuron Against Reproductive of Bactrocera carambolae Fruit Flies (Diptera: Tephritidae). The Journal of Experimental Life Sciences 6, 25–28 (2016).

41. Loof TG, et al. Coagulation, an ancestral serine protease cascade, exerts a novel function in early immune defense. Blood 118, 2589–2598 (2011).

42. Krem MM, Di Cera E. Evolution of enzyme cascades from embryonic development to blood coagulation. Trends Biochem Sci 27, 67–74 (2002).

